# Improved chemistry restraints for crystallographic refinement by integrating the Amber force field into Phenix

**DOI:** 10.1101/724567

**Authors:** Nigel W. Moriarty, Pawel A. Janowski, Jason M. Swails, Hai Nguyen, Jane S. Richardson, David A. Case, Paul D. Adams

## Abstract

The refinement of biomolecular crystallographic models relies on geometric restraints to help address the paucity of experimental data typical in these experiments. Limitations in these restraints can degrade the quality of the resulting atomic models. Here we present an integration of the full all-atom Amber molecular dynamics force field into Phenix crystallographic refinement, which enables a more complete modeling of biomolecular chemistry. The advantages of the force field include a carefully derived set of torsion angle potentials, an extensive and flexible set of atom types, Lennard-Jones treatment of non-bonded interactions and a full treatment of crystalline electrostatics. The new combined method was tested against conventional geometry restraints for over twenty-two thousand protein structures. Structures refined with the new method show substantially improved model quality. On average, Ramachandran and rotamer scores are somewhat better; clash scores and MolProbity scores are significantly improved; and the modelling of electrostatics leads to structures that exhibit more, and more correct, hydrogen bonds than those refined with traditional geometry restraints. We find in general that model improvements are greatest at lower resolutions, prompting plans to add the Amber target function to real-space refinement for use in electron cryo-microscopy. This work opens the door to the future development of more advanced applications such as Amber-based ensemble refinement, quantum mechanical representation of active sites and improved geometric restraints for simulated annealing.

**IMPORTANT:** this document contains embedded data - to preserve data integrity, please ensure where possible that the IUCr Word tools (available from http://journals.iucr.org/services/docxtemplate/) are installed when editing this document.

**Synopsis:** The full Amber force field has been integrated into Phenix as an alternative refinement target. With a slight loss in speed, it achieves improved stereochemistry, fewer steric clashes and better hydrogen bonds.

## 1. Introduction

Accurate structural knowledge lies at the heart of our understanding of the biomolecular function and interactions of proteins and nucleic acids. With close to 90% of structures in the Protein Data Bank (Berman *et al.*, 2000) solved via x-ray diffraction methods, crystallography is currently the pre-eminent method for determining biomolecular structure. Crystal structure refinement is a computational technique that plays a key role in post-experiment data interpretation. Refinement of atomic coordinates entails solving an optimization problem to minimize the residual difference between the experimental and model structure factor amplitudes (Jack & Levitt, 1978; Agarwal, 1978; Murshudov *et al.*, 1997). However, due to inherent experimental limitations and a typically low data to parameter ratio, the employment of additional restraints, commonly referred to as geometry or steric restraints, is key to successful structural refinement (Waser, 1963). These restraints, which can be thought of as a prior in the Bayesian sense, provide additional observations in the optimization target and reduce the danger of overfitting. Their use leads to higher quality, more chemically accurate models.

Most current refinement programs (Afonine *et al.*, 2012; Murshudov *et al.*, 2011; Sheldrick, 2008; Bricogne *et al.*, 2011) employ a set of covalent-geometry restraints first proposed by Engh & Huber in 1991 and later augmented and improved in 2001 (Engh & Huber, 1991, 2001). This set of restraints is based on a survey of accurate high-resolution small molecule crystal structures from the Cambridge Structural Database (Groom *et al.*, 2016) and includes restraints on interatomic bond lengths, bond angles and ω torsion angles. In addition, parameters are added to enforce proper chirality and planarity; multiple-minimum targets for backbone and side chain torsion angles; and repulsive terms to prevent steric overlap between atoms. Those terms are defined from small-molecule and high-resolution macromolecular crystal structure data and from interaction-specified van der Waals radii. They are very similar but not identical between refinement programs.

The Engh & Huber restraints function reasonably well, while the additional terms have been gradually improved, but a number of limitations have been identified over the years. Some of these limitations include: a lack of adjustability to differences in local conformation, protonation, and hydrogen bonding and to their changes during refinement; incomplete or inaccurate atom types and parameters for ligands, carbohydrates, and covalent modifications; use only of repulsive and not attractive steric terms; omission of explicit hydrogen atoms and their interactions; misleading targets resulting from experimental averaging artifacts; inaccurate dihedral restraints; and lack of awareness of electrostatic and quantum dispersive interactions with a consequent lack of accounting for hydrogen bonding cooperativity (Priestle, 2003; Touw & Vriend, 2010; Davis *et al.*, 2003; Moriarty *et al.*, 2014; Tronrud *et al.*, 2010).

Phenix (Liebschner *et al.*, 2019) includes a built-in system for defining ligand parameters (Moriarty *et al.*, 2009) that by default restrains the explicit hydrogen atoms at electron-cloud-center positions for X-ray and optionally at nuclear positions for neutron crystallography (Williams, Headd *et al.*, 2018). Addition of the Conformation Dependent Library (CDL) (Moriarty *et al.*, 2014), which makes backbone bond lengths and angles dependent on ϕ,ψ values, has improved the models obtained from refinement at all resolutions, and thus is the default in Phenix refinement (Moriarty *et al.*, 2016). Similarly, Phenix uses ribose-pucker and base-type dependent torsional restraints for RNA (Jain *et al*., 2015). For bond lengths and angles, protein side chains continue to use standard Engh & Huber restraints while RNA/DNA use early values (Gelbin *et al*., 1996; Parkinson *et al*., 1996) with a few modifications. This use of combined restraints is here designated CDL/E&H.

An alternative approach is the use of geometry restraints based on all-atom force fields used for molecular dynamics studies. This is not a novel idea. In fact, some of the earliest implementations of refinement programs employed molecular mechanics force fields (Jack & Levitt, 1978; Brünger *et al.*, 1987, 1989). However, at the time, restraints derived from coordinates of ideal fragments (Tronrud *et al.*, 1987; Hendrickson & Konnert, 1980) were found to provide better refinement results. The insufficiency of molecular mechanics-based restraints was mainly attributed to two factors: inaccurate representation of chemical space because of too few atom types, and biases in conformational sampling resulting from unshielded electrostatic interactions. Subsequently, however, the methods of molecular dynamics and corresponding force fields have seen significant development and improvement. Current force fields contain more atom types and are easily adjustable as needed. They are typically parameterized against accurate quantum mechanical calculations, not feasible just a few years ago, as well as using more representative experimental results. Significant methodological advances, such as the development of Particle Mesh Ewald (York *et al.*, 1993; Darden *et al.*, 1993) for accurate calculation of crystalline electrostatics and improved temperature and pressure control algorithms, have greatly increased accuracy. Modern force fields have been shown to agree well with experimental data (Zagrovic *et al.*, 2008; van Gunsteren *et al.*, 2008; Showalter & Brüschweiler, 2007; Grindon *et al.*, 2004; Bowman *et al.*, 2011), including crystal diffraction data (Cerutti *et al.*, 2009; Janowski *et al.*, 2013; Cerutti *et al.*, 2008; Liu *et al.*, 2015; Janowski *et al.*, 2015).

We have made it possible to use of the Amber molecular mechanics force field as an alternative source of geometry restraints to those of CDL/E&H. Here we present an integration of the Phenix software package for crystallographic refinement, *phenix.refine* (Afonine *et al.*, 2012) and the Amber software package (Case *et al.*, 2018) for molecular dynamics. We present results of paired refinements for 22,544 structures and compare Amber to traditional refinement in terms of model quality, chemical accuracy and agreement with experimental data, studied both for overall statistics and for representative individual examples. We also describe the implementation and discuss future directions.

## 2. Methods

### 2.1. Code preparation

The integration of the Amber code into *phenix.refine* uses a thin client. Amber provides a python API to its *sander* module, so that a simple “import sander” python command allows Phenix to obtain Amber energies and forces through a method call. At each step of coordinate refinement, Phenix expands the asymmetric unit coordinates to a full unit cell (as required by *sander*), combines energy gradients returned from Amber (in place of those from its internal geometric restraint routines) with gradients from the X-ray target function, and uses these forces to update the coordinates. Alternate conformers can take advantage of the “locally-enchanced-sampling” (LES) facility in *sander*: atoms in single-conformer regions interact with multiple-copy regions via the average energy of interaction, while different copies of the same group do not interact among themselves (Roitberg & Elber, 1991; Simmerling *et al.*, 1998).

The Amber files required are created by a preliminary *AmberPrep* program that takes a PDB file as input. It creates both a parameter-topology (*prmtop*) file used by Amber and a new PDB file containing a complete set of atoms (including hydrogens and any missing atoms) needed to do force field calculations. If requested, alternate conformers present in the input PDB file can be translated into *sander* LES format. For most situations, *AmberPrep* does not require the user to have any experience with Amber or with molecular mechanics; less-common situations (described in supporting information) require some familiarity with Amber. All the code required for both the *AmberPrep* and *phenix.refin*e steps is included in the current major release, 1.16-3549 and subsequent nightly builds of Phenix.

### 2.2. Structure selection and overall refinement protocol

To compare refinements using Amber against traditional refinements with CDL/E&H restraints, structures were selected from the Protein Data Bank (Burley *et al.*, 2019) using the following criteria. Entries must have untwinned experimental data available that are at least 90% complete. Each entry’s R_free_ was limited to a maximum of 35%, R_work_ to 30% and the **Δ**R (R_free_-R_work_) to a minimum of 1.5%. The lowest resolution was set at 3.5Å. Entries containing nucleic acids were excluded.

Coordinate and experimental data files were obtained directly from the Protein Data Bank (PDB) and inputs prepared via the automated *AmberPrep* program (see section 2.1 above). Entries containing complex ligands were included if the file preparation program *AmberPrep* was able to automatically generate and include the ligand geometry data; this generally excludes ligands containing covalent connections to the protein, or with metal atoms. Details of the internals of *AmberPrep* will be described elsewhere. Resolution bins (set at 0.1Å) with less than 10 refinement pairs were eliminated to reduce noise caused by limited statistics. Complete graphs are included in the supplemental material. The resulting 22,000+ structures had experimental data resolutions between 0.8Å and 3.6Å, with most of the structures in the 1.2-3.0 Å range (see figure 1).

**Figure 1.**
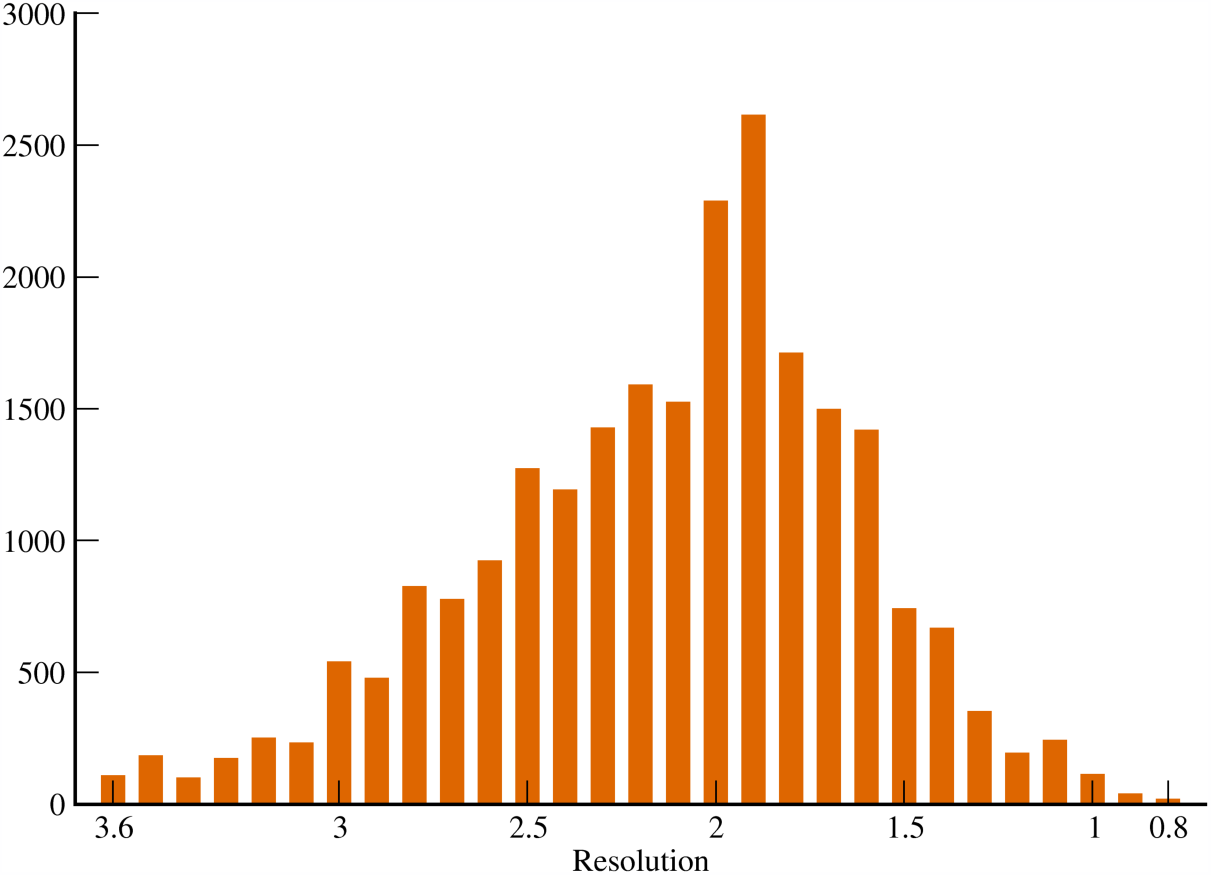
Distribution of refined structures across resolution bins.

Each model was then subjected to 10 macrocyles of refinement using the default strategy in *phenix.refine* for reciprocal space coordinate refinement, with the exception that real space refinement was turned off. By default, the first macrocycle uses a least-squares target function and the rest use maximum likelihood. Other options applied to both CDL/E&H and Amber refinements included optimization of the weight between the experimental data and the geometry restraints. This protocol was performed in parallel, once using CDL/E&H and once using Amber geometry restraints. In addition, Cβ pseudo-torsion restraints were not included in the restraints model. Explicit parameter settings are included in the supplemental material. Only one copy of each alternate conformation was considered initially (i.e. alternative location A). The final files are available by contacting the corresponding author.

The quality of the resulting models was assessed numerically using MolProbity (Williams, Headd *et al.*, 2018) available in Phenix (Adams *et al.*, 2010), by *cpptraj* (Roe & Cheatham, 2013) available in *AmberTools* (Case *et al.*, 2018) and by visual inspection with electron density and validation markup in KiNG (Chen *et al.*, 2009). All-atom dots for figure 10 were counted in Mage (Richardson & Richardson, 2001) and figures 5-9 were made in KiNG. To avoid typographical ambiguity, PDB codes are given here with lower case for all letters except L (e.g., 1nLs) (Moriarty, 2015).

**Figure 2.**
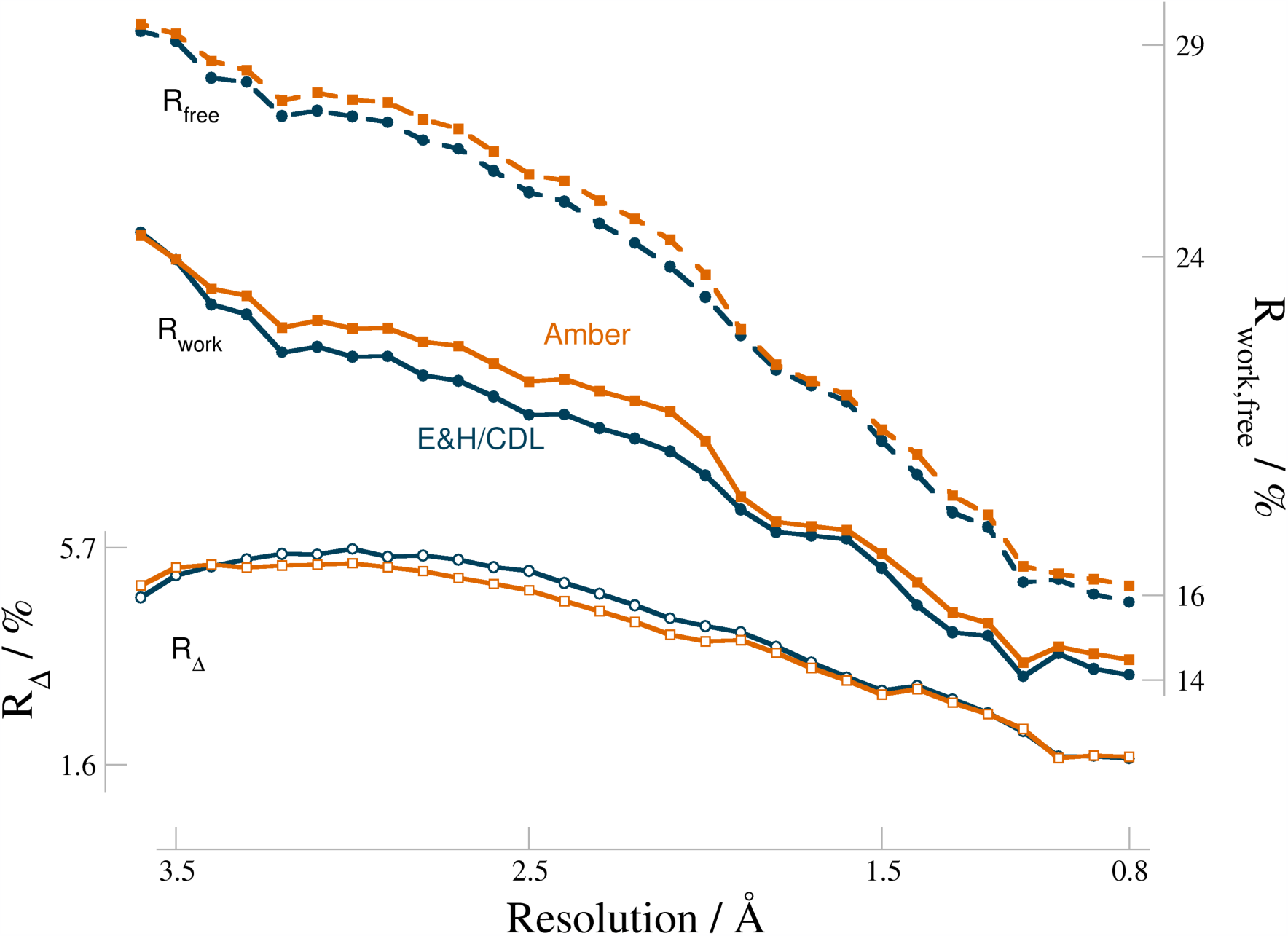
R-factors of optimized weight refinements and Rfree-Rwork (R_Δ_), versus resolution (values averaged in each resolution bin). Vertical axes are in % with R_Δ_ axis on the left. E&H/CDL values are plotted in dark blue and Amber in burnt orange.

**Figure 3.**
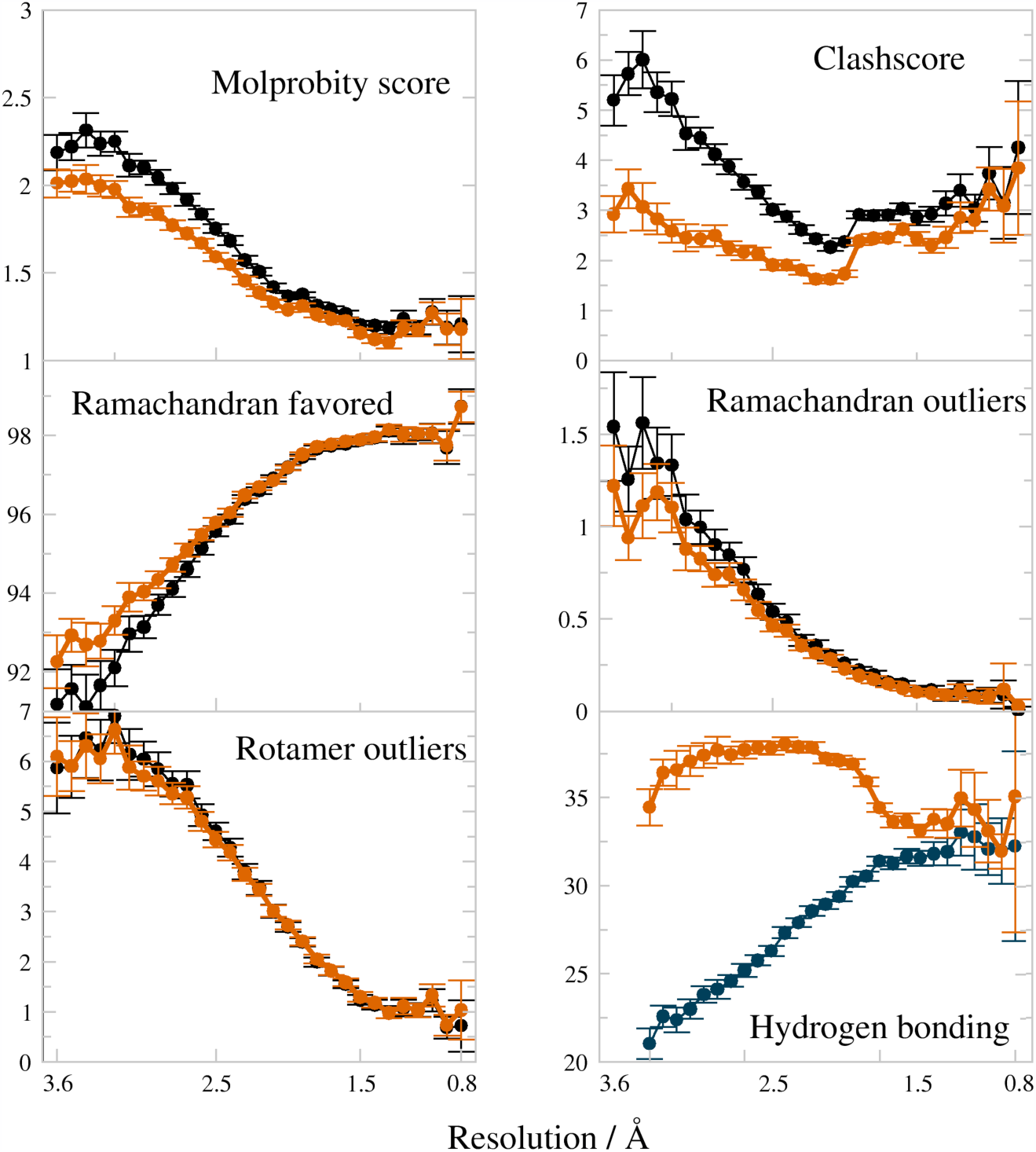
Comparison plots of model quality measures vs resolution, for Amber (burnt orange) vs CDL/E&H (dark blue) refinements with error bars depicting the 95% confidence level of the standard error of the mean. MolProbity score is a combination of all-atom clashscore, Ramachandran favored and rotamer outliers, weighted to approximate the expected score at the structure’s resolution. The hydrogen bond fraction is calculated using *cpptraj* per 1000 atoms in the model. For all 6 plots, Amber (burnt orange) differs in the better direction.

**Figure 4.**
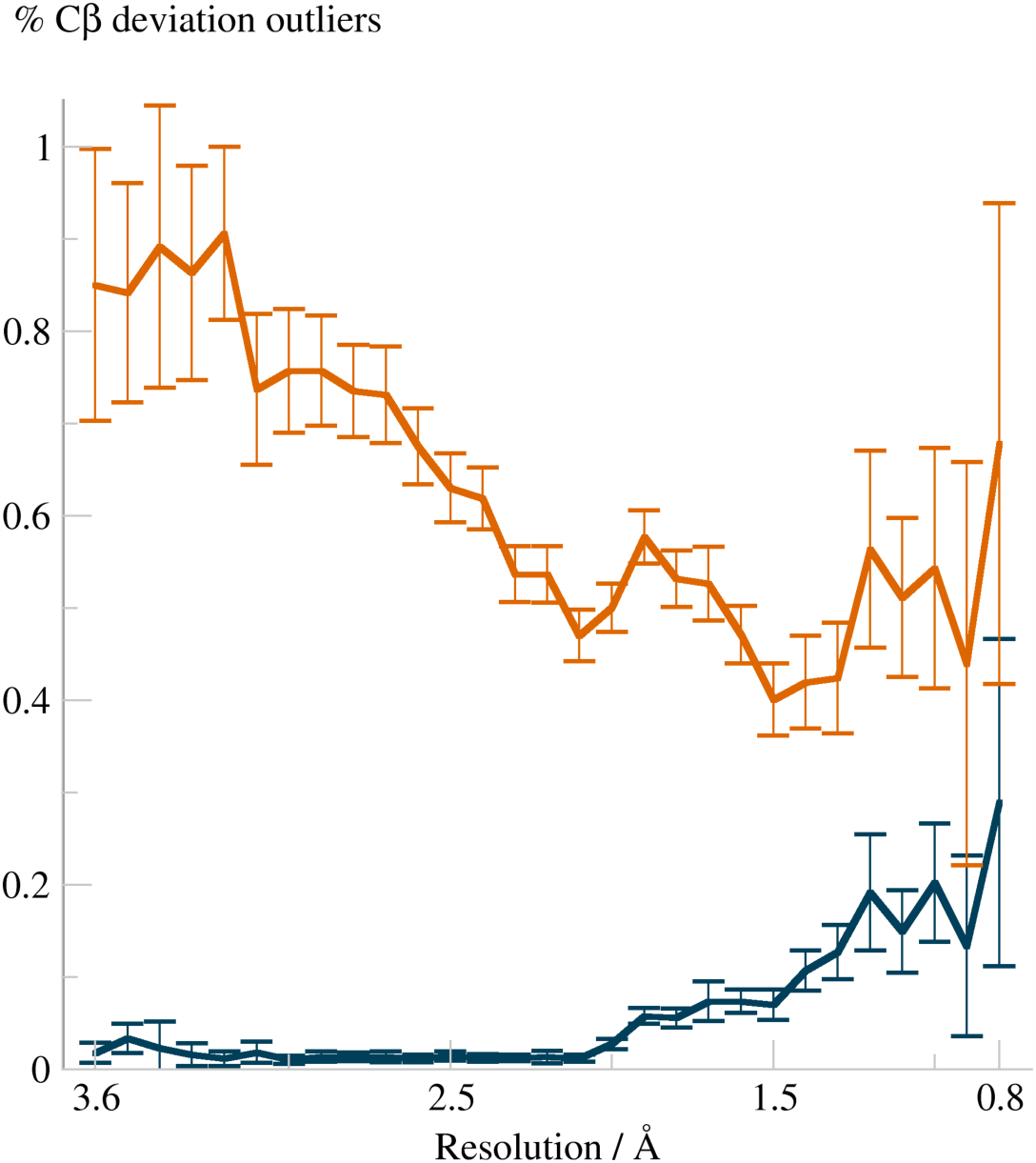
Fraction of Cβ deviations (in %) per Cβ atoms as a function of resolution, for the CDL/E&H (dark blue) and Amber (burnt orange) refinements. Values are averaged in each bin of resolution, with the error bars showing the 95% confidence level of the standard error of the mean.

**Figure 5.**
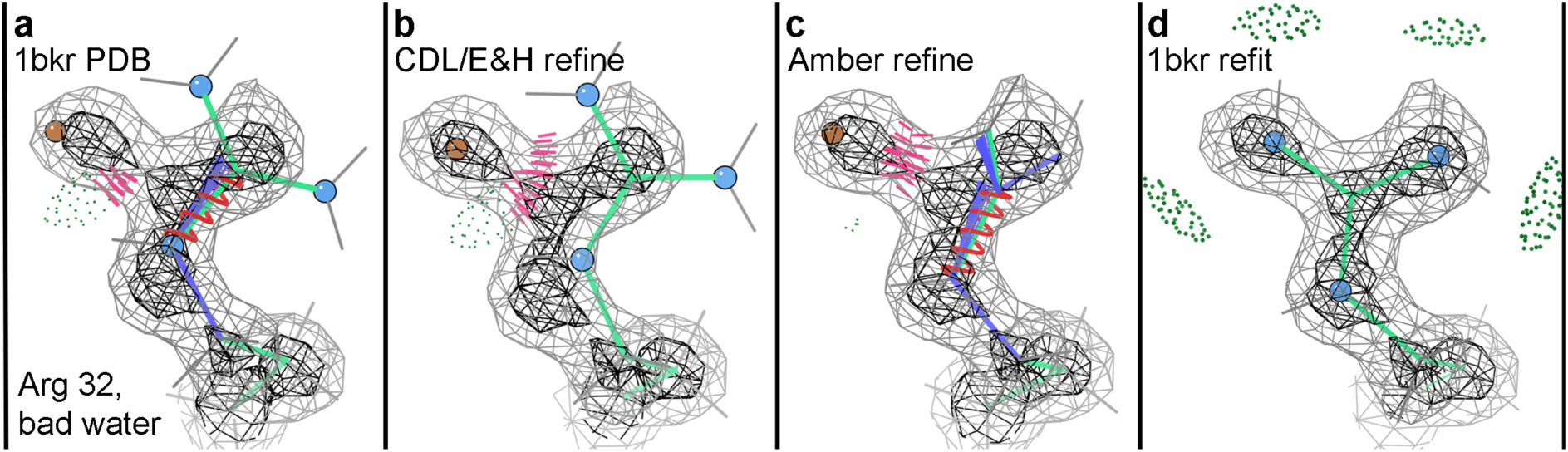
Differing responses of CDL/E&H versus Amber refinement to the misfitting of a water into what should be a side chain N atom in an Arginine. Neither result here is acceptable, but if the incorrect water is deleted (panel d), both methods do a very good job of moving the guanidinium correctly back into its density. MolProbity markup for figures 5-10: clusters of hotpink spikes for clashes, pillows of green dots for H-bonds, red or blue springs or fans for larger or smaller bond length or angle outliers, magenta spheres for Cβ deviations, gold sidechains for rotamer outliers, green Cα-Cα lines for Ramachandran outliers, and magenta lines along the CO-CO dihedral for CaBLAM outliers. Relevantly moving O or N atoms are emphasized with red or blue balls.

**Figure 6.**
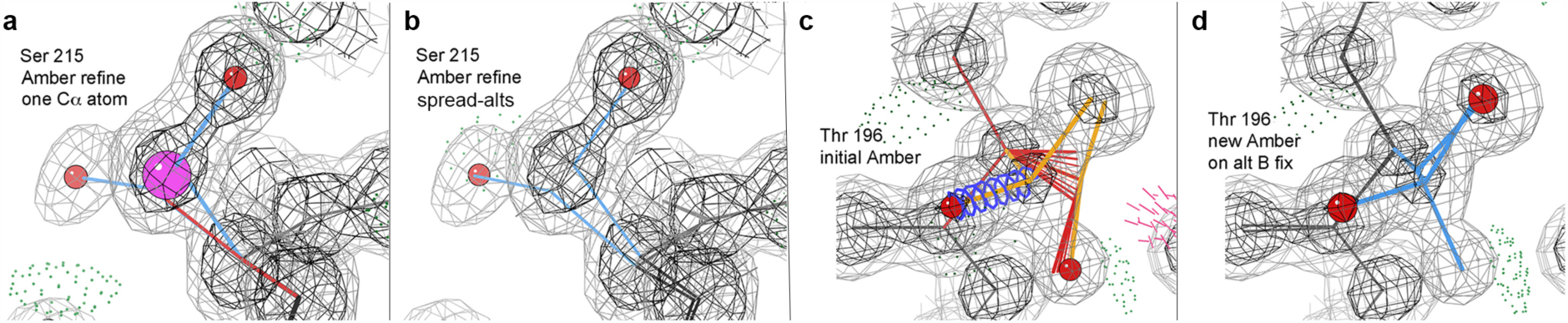
At high resolution, Cβ deviation outliers are most often due to problems with alternate conformations. a) Amber refinement using the original Ser 215 alternates in PDB file 1nLs, which have widely separated positions for Cβ but only a single Cα atom. b) Amber refinement after the definition of alternates has been spread to include the Cα and both adjoining peptides. c) Amber refinement of the original Thr 196 of 1nLs, where alternate B had been fit backward; there is bad covalent geometry and a huge Cβd of 0.88Å (ball not shown). d) Good Amber result after altB was refit in the correct rotamer, so that all atoms match the density.

**Figure 7.**
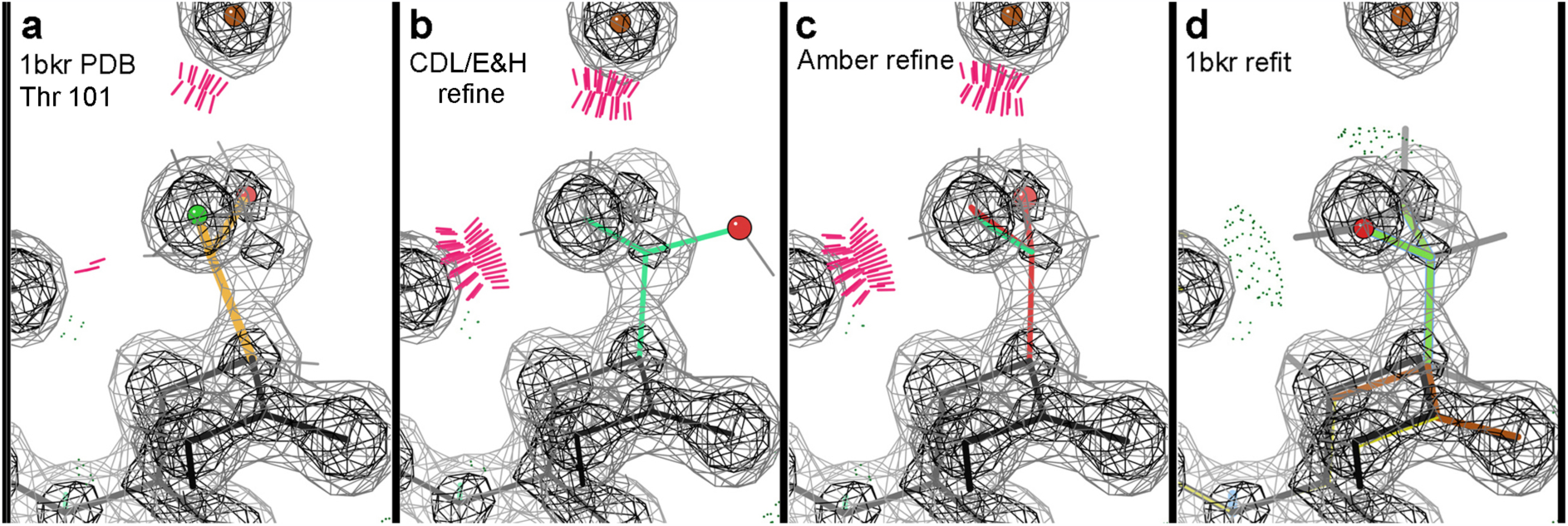
Unacceptable ways to get rid of a Cβ deviation without fixing the actual problem. a) 1bkr Thr 101 as deposited, with a huge Cβd of 0.63Å (not shown as a ball because it obscures the side chain), clashes, a rotamer outlier, the heavier Oγ branch in the lower electron-density peak and the Cβ out of density -- all caused by modeling the side chain χ1 180° backwards. b) CDL/E&H makes the geometry perfect but puts the Oγ far out of density. c) Amber gets all 3 side chain atoms into peaks by making the chirality at Cβ incorrect. d) A refit in the correct rotamer replaces clashes with H-bonds, has no outliers and puts each atom into its correct density peak.

**Figure 8.**
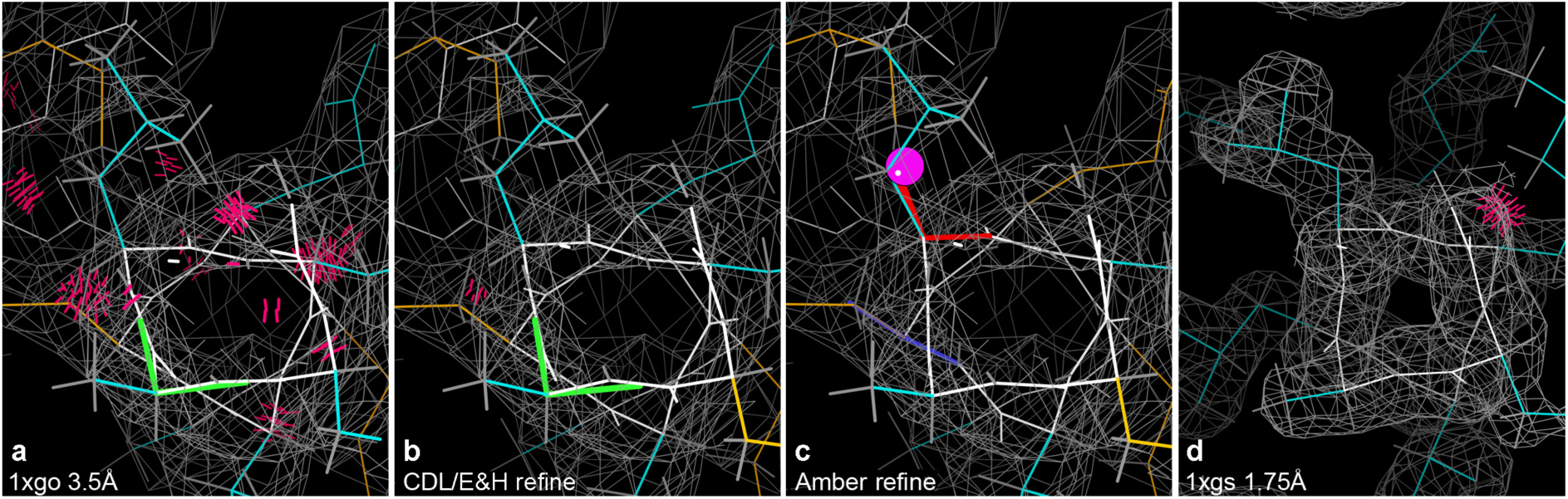
A Cβ deviation in the Amber results at 3.5Å, but not for either the original or the CDL results. a) 1xgo Leu 253 on a quite distorted helix, with many clashes and a Ramachandran outlier; the Leu rotamer is incorrect, as shown by the 1xgs structure at 1.75Å. b) CDL/E&H refinement fixes the clashes, but not the rotamer or Ramachandran outliers or the helix distortion. c) Amber refinement fixes the clashes and the Ramachandran outlier, flags the incorrect Leu rotamer with a Cβd outlier and moves the helix conformation closer to ideal. d) Leu 253 in 1xgs at 1.75Å, with a clearly correct rotamer on an ideal helix and no outliers besides one clash.

**Figure 9.**
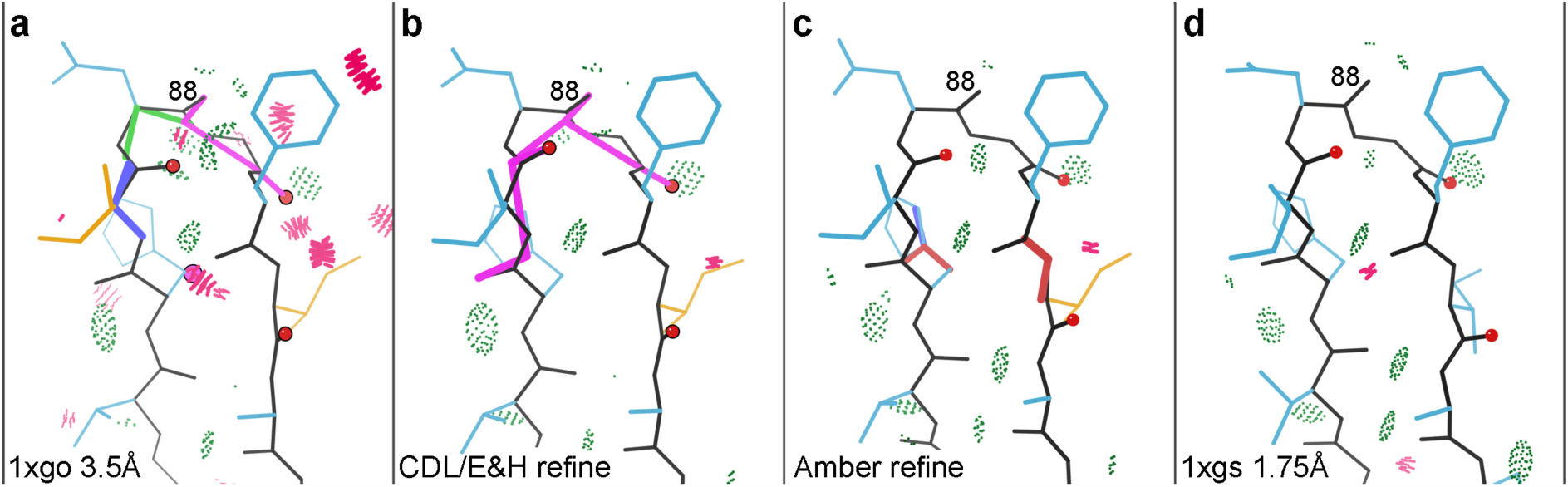
Two misoriented peptides in 1xgo, flagged by Ramachandran and CaBLAM outliers (magenta outlines on the CO virtual dihedrals). a) Residues 86-91 in the deposited 1xgo structure. b) CDL/E&H result, with unchanged conformation and outliers. c) Amber result, with several peptide orientations changed by modest amounts (red balls on CO), removing the backbone outliers and very closely matching the conformation for 1xgs shown in panel d.

**Figure 10.**
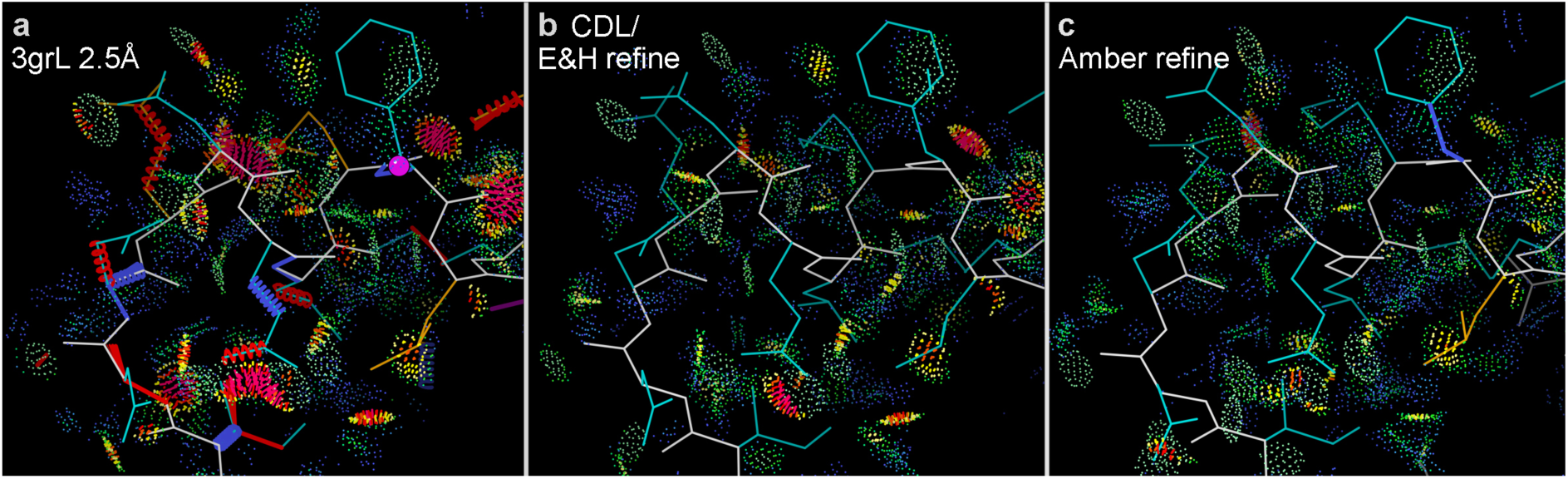
Amber refinement produces better H-bonds and van der Waals contacts as well as removing somewhat more steric clashes. a) The Asn 182 helix-cap region in PDB file 3g8L at 2.5Å, with numerous clashes and other outliers. b) CDL/E&H refinement makes large improvements, removing most clashes and all other outliers. c) Amber refinement does even better, removing all clashes and most small overlaps (yellow) and optimizing to produce more H-bonds and favorable van der Waals contacts (green and blue dots).

### 2.3. Weight factor details

The target function optimized in *phenix.refine* reciprocal space atomic coordinate refinement is of the general form:

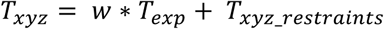

where all the terms are functions of the atomic coordinates, *T*_*xyz*_ is the target residual to be minimized, *T*_*exp*_ is a residual between the observed and model structure factors and quantifies agreement with experimental data, *T*_*xyz_restraints*_ is the residual of agreement with the geometry restraints and *w* is a scale factor that modulates the relative weight between the experimental and the geometry restraint terms. In traditional refinement *T*_*xyz_restraints*_ is calculated using the set of CDL/E&H restraints:

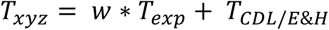

To implement Phenix-Amber we substitute this term with the potential energy calculated using the Amber force field:

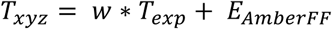

where the Amber term is intentionally represented now by an *E* to emphasize that we directly incorporate the full potential energy function calculated in Amber using the ff14SB (Maier *et al.*, 2015) force field.

In a standard default Phenix refinement, the weight, *w*, is a combination of a value based on the ratio of gradient norms (Brünger *et al.*, 1989; Adams *et al.*, 1997) and a scaling factor that defaults to ½. This initial weight can be optimized using a procedure described previously (Afonine *et al.*, 2011). This procedure uses the results of ten refinements with a selection of weights, considering the bond and angle rmsd, the R-factors and validation statistics to determine the best weight for the specific refinement at each of the ten macrocycles. The same procedure was used to estimate an optimal weight for the Phenix-Amber refinements. (If faster fixed-weight refinements are desired, we have found that a scaling factor of 0.2, rather than 0.5, scales the Amber gradients to be close to those from the CDL/E&H restraints, allowing the simpler, default, weighting scheme in *phenix.refine* to be used.)

## 3. Results

### 3.1. Full-dataset score comparisons

On average, the Phenix-Amber combination produced slightly higher R-work and R-free (figure 2) but higher quality models (figure 3). The increase in R-factors is most pronounced in the 1.8–2.8Å range. This is a result of the weight optimisation procedure having different limits for optimal weight in this resolution range. The increase was less for R-free than R-work such that the R-delta is less for refinements using Amber gradients. The uncertainty in the R-free for 95% of refinements calculated using equation 13 of (Tickle *et al.*, 2000) is less than 0.032. At 2Å resolution, this equates to an uncertainty of 0.7% which is approximately the same as the difference in the average R-free values of 23.0% and 23.6% for Phenix and Phenix-Amber, respectively.

The Phenix-Amber refinements exhibited improved (lower) MolProbity scores and contained fewer clashes between atoms. Plots show the mean of the values in the 0.1Å resolution bin as well as the 95% confidence level of the standard error of the mean (SEM). MolProbity clashscores are particularly striking: for refinement using CDL/E&H restraints, clashscores steadily increase as resolution worsens, often resulting in very high numbers of steric clashes. On the other hand, the mean clash-score with Amber restraints appears to be nearly independent of resolution and remains consistent at about 2.5 clashes per 1000 atoms across all resolution bins. The SEM range is non-overlapping for worse than 1Å indicating that the Amber force field is producing better geometries at mid to low resolution. There are more favored Ramachandran points (backbone ϕ,ψ) and fewer Ramachandran outliers for the Phenix-Amber refinements. This difference is most marked for resolutions worse than 2Å. Phenix-Amber refinement also improves (lowers) the number of rotamer outliers but doesn’t differentiate via the SEM, and increases the proportion of hydrogen bonds. While the rotamer outlier results remain similar, the hydrogen bonding results have a large difference at worse than 2Å resulting in nearly double the bonds near 3Å. Common to all the plots is a change near 2Å, where the weight optimisation procedure common to both CDL/E&H and Amber refinement loosens the weight on geometry restraints somewhat, to allow more deviations at resolutions where the data is capable of unambiguously showing them. Bond and angle rmsd comparisons are less pertinent as the force fields do not have ideal values for parameterisations and comparing the Phenix-Amber bonds and angles to the CDL/E&H values is not a universal metric. The curious can see the plots in figure S1. Overall, improvement with Amber is substantial in the lower resolution refinements.

One validation metric that is worse for Phenix-Amber refinements is the number of outliers of the Cβ positions. Both the mean and the SEM show clear differentiation. The Cβ deviation is the distance between the modelled Cβ and an ideal Cβ, that is a combined measure of distortion in the tetrahedron around the Cα atom. The ideal position is calculated by averaging the N-C-Cα-Cβ and C-N-Cα-Cβ improper dihedrals and correcting bond length, which allows for the effect of a non-ideal tau angle (Lovell *et al.*, 2003). With traditional E&H restraints the Cβd is quite robustly sensitive to incompatibility between how the backbone and side chain conformations have been modelled. For CDL/E&H refinements, however, the percentage of Cβd outliers (>0.25Å) is negligible for low and mid resolutions, only increasing to 0.2% at higher resolutions (see figure 4). This is in line with the CDL/E&H providing tight geometrical restraints out to Cβ at most resolutions, but loosened somewhat at better than 2Å resolution where there is enough experimental information to move an angle away from ideal. Note that explicit Cβ restraints were turned off for all Phenix refinements and that the Amber force field does not have an explicit Cβ term; however, if all angles around the Cα are kept ideal then the Cβ position will also be ideal even if it is incorrectly positioned in the structure. The following section analyses specific local examples where output structures show differences for either the positive or the negative trends seen in the overall comparisons, in order to understand their nature, causes and meaning across resolution ranges.

### 3.2. Examination of individual examples

As noted above, in comparison with the CDL/E&H restraint refinements, the Phenix-Amber refinements have much higher percentages of Cβ deviation outliers, increasing at the low-resolution end to more than 1% of Cβ atoms. Amber refinement also has more bond length and angle outliers. The following examines a sample of cases at high, mid and lower resolutions to understand the starting-model characteristics and refinement behaviour that produce these differences.

#### 3.2.1. High resolution: waters, alternates, Cβd outliers and atoms in the wrong peak

In the high-resolution range (better than 1.7Å), it appears that the commonest problems not easily correctable by refinement are caused either by modeling the wrong atom into a density peak or by incorrect modeling, labeling, or truncation of alternate conformations. Such problems are usually flagged in validation either by all-atom clashes, by Cβ deviations and sometimes by bad bond lengths and angles. (For the high-resolution examples described here, we use the LES procedure outlined above to model alternative conformers in the Phenix-Amber refinements.)

Figure 5a shows a case where a water molecule had been modeled in an electron density peak that should really be a nitrogen atom of the Arg guanidinium. CDL/E&H refinement (figure 5b) corrected the bad geometry at the cost of moving the guanidinium even further out of density; Amber refinement changed the guanidinium orientation but made no overall improvement (figure 5c); all three versions have a bad clash. If the water were deleted, then either refinement method would undoubtedly do an excellent job (figure 5d). This type of problem is absent at low resolution where waters are not modeled but occurs quite often at both high and mid resolution, for other branched side chains, for Ile Cδ (for example, 3js8 195) and even occasionally for Trp (e.g. 1qw9 B170).

Cβ deviation outliers (≥0.25Å) are often produced by side chain alternates with quite different Cβ positions but no associated alternates defined along the backbone. Since the tetrahedron around Cα should be nearly ideal, that treatment almost guarantees bad geometry. The rather simple solution, implemented in Phenix, is to define alternates for all atoms until the i+1 and i-1 Cα atoms – as in the “backrub” motion; (Davis *et al.*, 2006). PDB codes 1dy5, 1gwe and 1nLs each have a number of such cases. Figure 6a,b show 1nLs Ser 215, initially with an outlier Cβd, 0.49Å distance between the two Cβ atoms and a single Cα. CDL/E&H refinement pulls the Cβ atoms to be only 0.23Å apart, avoiding a Cβd with only slightly worse fit to the density; Amber reduces the Cβd only slightly, but it does keep this flag of an underlying problem. When alternates are defined for the backbone peptides, both systems improve.

Worse cases occur where one or both alternates have been fit incorrectly as well as not being expanded along the backbone appropriately. Figure 6c shows Thr 196, with a huge Cβd of 0.88Å (not shown) and very poor geometry, because altB was fit incorrectly (just as a shift of altA rather than as a new rotamer). This time even CDL/E&H refinement produces a Cβd outlier, but smaller than for Amber. Figure 6d shows the excellent Amber result after the misfit of altB was approximately corrected.

#### 3.2.2. Mid resolution: backward side chains and rare conformations

An even commoner case at both high and mid resolutions where the wrong atom is fit into a density peak is a backward-fit Cβ-branched residue, well illustrated by a very clear Thr example in 1bkr at 1.1Å (figure 7a). Thr 101 is a rotamer outlier (gold) on a regular α-helix with a Cβd of 0.63Å. The deposited Thr 101 also has a bond-angle deviation of 13.5σ; clashes at the Cγ methyl; its Cβ is out of density; Oγ is in the lower peak; and Cγ is in the higher peak. It is shown in figure 7 with 1.6s and 4s 2mF_o_-DF_c_ contours (but without Cβ deviation and angle markups for clarity). This mistake was not obvious because anisotropic B’s were used too early in the modeling resulting in the Thr Cβ being refined to a 6:1 aniso-axis ratio that covered both the modeled atom and the real position. The figures show the density as calculated with isotropic B factors.

Given this difficult problem for automated refinement, each of the two target functions reacts very differently. Both refinements still have the Cγ methyl clashing with a helix backbone CO in good density, very diagnostic of a problem with the Cγ. It is indeed the wrong atom to have in that peak, as shown also by the relative peak heights. The CDL/E&H refinement (figure 7b) achieves tight geometry and a good rotamer, moving the Cβ into its correct density peak, but pays the price for not correcting the underlying problem by swinging the Oγ out of density. The Amber refinement (figure 7c) achieves an atom in each of the three side chain density peaks, but pays the price for not correcting the underlying problem by having the wrong chirality at the Cβ atom. It still also has bond-angle outliers, which may be a sign of unconverged refinement.

The original PDB entry, the CDL/E&H refinement and the Amber refinement structures for Thr 101 are all very badly wrong, but each in an entirely different way. The deposited model, 1bkr, looks very poor by traditional model validation, but has a misleadingly good density correlation, given the extremely anisotropic Cβ B-factor. The CDL/E&H output looks extremely good on traditional validation except for the clashes and would show a lowered but still reasonable density correlation; however, it is the most obviously wrong upon manual inspection. The Amber output has clashes and currently has modest bond-angle outliers, but it fits the density very closely making it difficult to identify as incorrect by visual inspection. The problem could be recognized automatically by a simple chirality check. Shown in figure 7d, Thr 101 was rebuilt quickly in KiNG, with the **p** rotamer and a small backrub motion. Either Phenix-CDL/E&H or Phenix-Amber refinement would do a very good job from such a rough refit with the correct atoms near the right places.

At mid resolution, there are also other rotamers and backbone conformations fit into the wrong local minimum and thus difficult to correct by minimization refinement methods, but not always flagged by Cβ deviations or other outliers. Some of these, such as *cis*-nonPro peptides (Williams, Videau *et al.*, 2018) or very rare rotamers (Hintze *et al.*, 2016) can be avoided by considering their highly unfavorable prior probabilities. Others would require explicit sampling of the multiple minima.

#### 3.2.3. Lower resolution: peptide orientations with CaBLAM and Cβd outliers

At low resolution (2.5–4Å), no waters or alternates are modeled. All other problems continue, but an additional set of common local misfittings occur because the broad electron density is compatible with significantly different models. 1xgo at 3.5Å is an excellent case for testing in this range, because it was solved independently from the 1.75Å 1xgs structure – the same molecule in a different space group. CDL/E&H refinement shows no Cβd outliers, but Amber refinement has six. Comparison with 1xgs shows that each of the Cβd residues has either the side chain or the backbone or both in an incorrect local-minimum conformation uncorrectable by minimization refinement methods (Richardson & Richardson, 2018). For example, figure 8 shows Leu 253 on a helix, with a Cβd from Amber (panel c) and the different, correct 1xgs Leu rotamer in panel d. Those Cβd outliers are thus a feature, not a bug, in Amber: they serve their designed validation function of flagging genuine fitting problems. However, the lack of Cβd outliers in the CDL/E&H refinement is also not a defect, because the tight CDL/E&H geometry is on average quite useful at low resolution.

The 1xgo-vs-1xgs comparison also illustrates many of the ways in which Amber refinement is superior at low resolution. In figure 8, Amber corrects a Ramachandran outlier in the helix and shows a helix backbone shape much closer to the ideal geometry of 1xgs than either the deposited or the CDL/E&H versions.

Since the backbone CO direction cannot be seen at low resolution, the commonest local misfitting is a misoriented peptide (Richardson *et al.*, 2018). Those can be flagged by the new MolProbity validation called CaBLAM, which tests whether adjacent CO directions are compatible with the local Cα backbone conformation (Williams, Headd *et al.*, 2018). Ten such cases were identified in 1xgo, for isolated single or double CaBLAM outliers surrounded by correct structure as judged in1xgs. For six of those 10 cases, neither CDL/E&H nor Amber refinement corrected the problem: His62, Thr70, Gly163, Gly193, Ala217, Glu286 (see stereo figure S2). In two cases CDL/E&H had fewer other outliers than Amber, but did not actually reorient the CO: for Gly193 and for the Gly163 case shown in figure S3. In three of the 10 cases Amber did a complete fix, while CDL/E&H did not improve (Asp88, Gly125, Pro266). For example, in figure 9, 1xgo residues 86-91 (panel a) have a CaBLAM outlier (magenta lines), uncorrected by CDL/E&H refinement (panel b). But Amber refinement (panel c) manages to shift several CO orientations by modest amounts (red balls), enough to fix the CaBLAM outliers and match extremely closely the better backbone conformation of 1xgs (panel d). The Gly 125 example is shown in figure S4. Finally, in one especially interesting case (Lys22) Amber turned the CO about halfway up to where it should be, while CDL/E&H made no improvement. The Amber model still has geometry outliers and further runs moved most of the way up and removed those outliers, showing that Amber refinement had not yet fully converged in 10 macrocycles (see Supplement text and figure S5).

Amber refinement is especially good at optimizing hydrogen-aware all-atom sterics, as calculated by the Probe program (Word, Lovell, LaBean *et al.*, 1999) with H atoms added and optimized by Reduce (Word, Lovell, Richardson *et al.*, 1999). This is illustrated in figure 10 for 3g8L at 2.5Å resolution. The deposited structure of the Asn 182 helix N-cap region, which has many outliers of all kinds (panel a), is improved a great deal by CDL/E&H refinement (panel b). However, the Amber refinement (panel c) is noticeably better, with more H-bonds and better van der Waals contacts as well as fewer clashes. These improvements are plotted quantitatively in figure 11, as measured by a drop in unfavorable clash spikes (red) and small overlaps (orange), with an increase in favorable H-bonds (green) and van der Waals contacts (blue).

**Figure 11.**
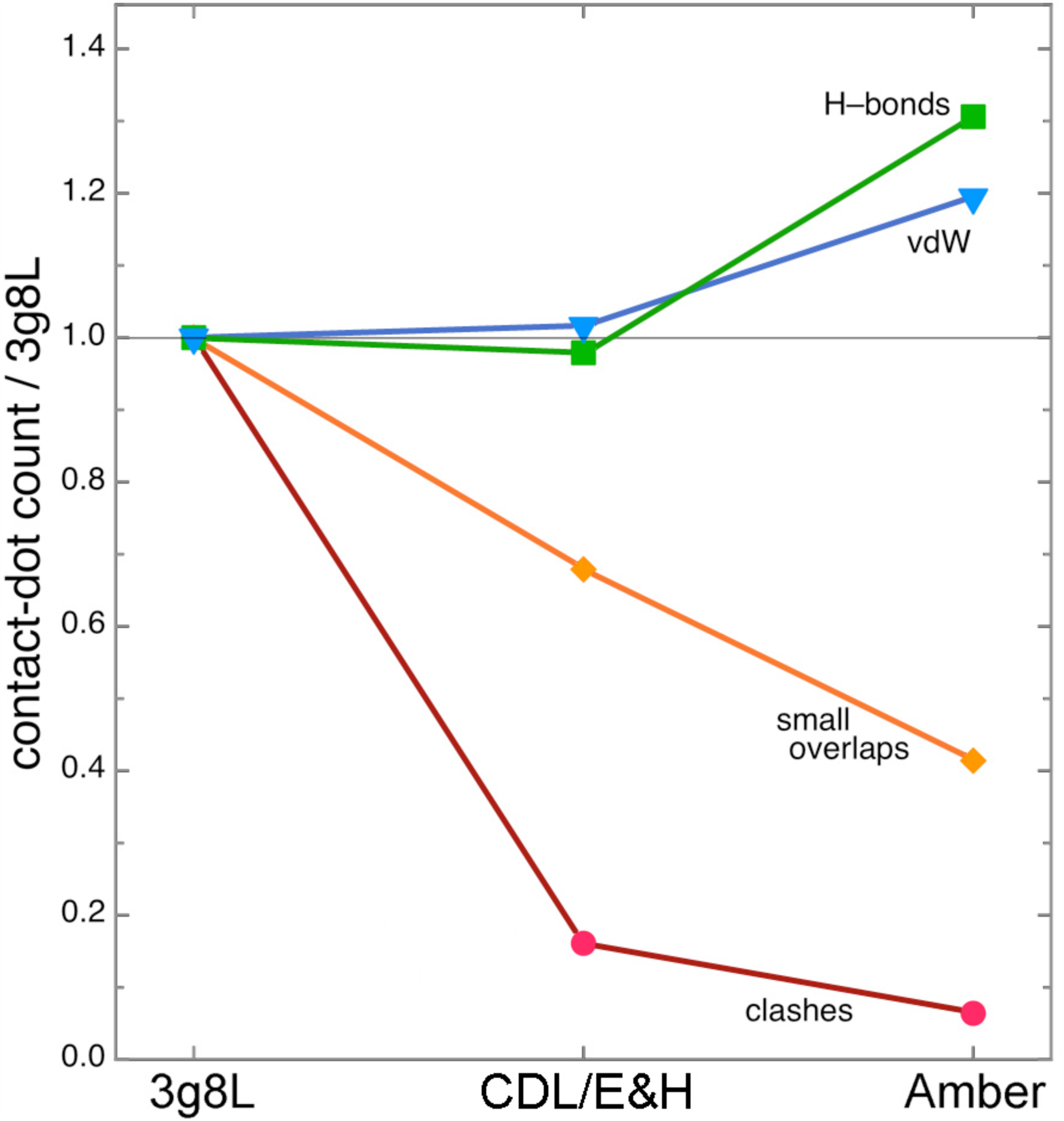
CDL/E&H versus Amber improvements in steric contacts for the 3g8L helix-cap, quantified by all-atom contact dot or spike counts measured in Mage (Richardson 2001), normalized relative to the counts in the deposited 3g8L structure. Amber changes farthest, in the right direction, for all four contact types.

## 4. Discussion

The idea of including molecular mechanics force fields into crystallographic refinements is not a new one, with precedents dating back to early work by (Jack & Levitt, 1978) and the XPLOR program (Brünger & Karplus, 1991) developed in the 1980’s. The notion that a force field could (at least in principle) encode “prior knowledge” about protein structure continues to have a strong appeal and efforts to replace conventional “geometric restraints”, which are very local and uncorrelated, with a more global assessment of structural quality have been explored repeatedly (e. g., Moulinier *et al.*, 2003; Schnieders *et al.*, 2009). Distinguishing features of the current implementation include automatic preparation of force fields for many types of biomolecules, ligands and solvent components as well as close integration with Phenix, a mature and widely used platform for refinement. This has enabled parallel refinements on more than 22,000 protein entries in the PDB and allows crystallographers to test these ideas on their own systems by simply adding flags to an existing *phenix.refine* command line or adding the same information via the Phenix GUI. Indeed, we expect most users to “turn on” Amber restraints after having carried out a more conventional refinement to judge for themselves the significance and correctness of structural differences that arise. As noted in Section 3.2, an Amber refinement will often flag residues that need manual refitting in ways complementary to the cues provided by more conventional refinement.

The results presented here show that structures with improved local quality (as monitored by MolProbity criteria and hydrogen bond analysis) can be obtained by simple energy minimization, with minimal degradation in agreement with experimental structure factors and with no changes to a current-generation protein force field. Nevertheless, one should keep in mind that the Amber-refined structures obtained here are not very different from those found with more conventional refinement. Both methods require that most local misfittings to be corrected in advance. The hope is that either sampling of explicit alternatives or else optimization using more aggressive conformational search, such as with simulated annealing or torsion-angle dynamics, may find the correct low-energy structures with good agreement with experimental data.

It is likely that further exploration of relative weights between “X-ray” and “energy” terms (beyond the existing and heuristic weight-optimization procedure employed here) and even within the energy terms, will become important. In principle, maximizing the joint probability arising from “prior knowledge” (using a Bolztmann distribution, exp(-E_AmberFF_/k_B_T) for some effective temperature) and a maximum likelihood target function (based on a given model and the observed data) is an attractive approach that effectively establishes an appropriate relative weighting. More study will be needed to see how well this works in practice, especially in light of the inevitable limitations of current force fields.

The integration of Amber’s force field into the Phenix software for crystallography also paves the way for the development of more sophisticated applications. The force field can accommodate alternate conformers by using the locally enhanced sampling (LES) approach (Roitberg & Elber, 1991; Simmerling *et al.*, 1998); a few examples are discussed here whilst details will be presented elsewhere. Ensemble refinement (Burnley *et al.*, 2012) could now be performed using a full molecular dynamics force field, thus avoiding poor quality individual models in the ensemble. Similarly, simulated annealing could now be performed with an improved physics-based potential. Extension of the ideas presented to real-space refinement within Phenix is underway, opening a path to new applications to cryo-EM and low-resolution X-ray structures. These developments would all contribute significantly to the future of macromolecular crystallography, reinforcing the transition from a single static-structure-dominated view of crystals to one where dynamics and structural ensembles play a central important role in describing molecular function (Furnham *et al.*, 2006; van den Bedem & Fraser, 2015; Wall *et al.*, 2014).

## 5. Conclusions

We have presented refinement results obtained by integrating Phenix with the Amber software package for molecular dynamics. Our refinements of over 22,000 crystal structures show that refinement using Amber’s all atom molecular mechanics force field outperforms CDL/E&H restraint refinement in many respects. An overwhelming majority of Amber-refined models display notably improved model quality. The improvement is seen across most indicators of model quality including clashes between atoms, side chain rotamers and peptide backbone torsion angles. In particular, Phenix-Amber consistently outperforms standard Phenix refinement in clashscore, number of hydrogen bonds per 1000 atoms and MolProbity score. It also consistently outperforms standard refinement for Ramachandran and rotamer statistics at low resolutions and obtains approximately equal results at high (better than 2.0Å) resolutions. Amber does run somewhat more slowly (generally 20-40% longer) and may take more cycles for a particular local conformation to converge completely if it is making a large local change (see text for supplementary figure S5). It should be noted that standard refinement consistently outperforms Phenix-Amber in eliminating Cβ deviation and other covalent-geometry outliers across all resolutions, but in most cases the Amber outliers serve to flag a real problem in the model.

As the quality of experimental data decreases with resolution, the improvement in model quality obtained by using Amber, as opposed to CDL/E&H restraints, increases. This improvement is especially striking in the case of clashscores, which appear to be nearly independent of experimental data resolution for Amber refinements. Additional improvement is seen in the modelling of electrostatic interactions, H-bonds and van der Waals contacts, which are currently ignored by conventional restraints. Improving lower-resolution structures is very important, since they include a large fraction of the most exciting and biologically important current structures such as the protein/nucleic acid complexes of big, dynamic molecular machines.

No minimization refinement method, including CDL/E&H and Amber, can in general correct local misfittings that were modeled in an incorrect local-minimum conformation, especially at relatively high resolutions. At lower resolution where the barriers are softer, Amber sometimes can manage such a change, while CDL/E&H still does not. It is, therefore, important and highly recommended that validation flags be consulted for the initial model and as many as feasible of the worst cases be fixed, before starting the cycles of automated refinement with either target.

### Software distribution

Amber was implemented in *phenix.refine* and is available in the 1.16-3549 version of Phenix and later. Instructions for using the *phenix.refine* Amber implementation are available in the version-specific documentation available with the distribution. The Amber codes are included in the Phenix distribution under the terms of the GNU lesser general public license (LGPL).

## Acknowledgements

JSR thanks David Richardson for help with some aspects of the individual-example analyses. The content is solely the responsibility of the authors and does not necessarily represent the official views of the National Institutes of Health, NIGMS, or DOE.

## Supporting information

### S1. Preparation of structures for Phenix-Amber refinement

The *AmberPrep* program prepares the files needed for the subsequent refinement step. For components (typically ligands) that are not standard amino acids, nucleotides, solvent or monatomic ions, the eLBOW routines (Moriarty *et al.*, 2009) are used to add hydrogen atoms and determine the most likely protonation and tautomeric states. These three-dimensional structures are then used in the standard way in Amber’s antechamber tool (Wang *et al.*, 2006) to assign charges and atom types using version 2.11 of the general Amber force field (GAFF) (Wang *et al.*, 2004). Proteins are modeled using the ff14SB force field (Maier *et al.*, 2015), water and related ions with the TIP3P model and associated parameters for ions (Jorgensen *et al.*, 1983; Joung & Cheatham, 2009).

This procedure will fail for ligands containing metal ions (since the GAFF force field currently only deals with organic moieties), and also for ligands that have covalent connections to the protein. For each of these cases, users familiar with the Amber software can build the needed component libraries using other Amber-based tools. Because such efforts are not yet fully automated, structures with metal-containing ligands or covalent connections were left out of the current caclulations.

After these component libraries are prepared, the coordinates of the system are expanded to a full unit cell, and Amber’s *tleap* program is used to construct topology and coordinate files in Amber format. Disulfide bonds and gaps in the sequence are identified and properly processed. A model file in PDB format for the asymmetric unit (for use by Phenix) is also created that contains any added hydrogen atoms or missing atoms; any atomic displacement parameters (ADPs) from the input PDB file are copied to this file; hydrogen atoms are assigned isotropic B-factors that match the heavy atoms to which they are bonded. For the main statistical analysis, only the most populated alternate conformer was selected, and assigned unit occupancy. As discussed in the text, for a selected set of structures, we also used an option in the code to include all alternate conformers present in the input PDB file.

During refinement, *phenix.refine* sees only a single asymmetric unit, as usual. At each step, when Amber restraints are required, these coordinates are expanded to a full unit cell, the Amber force field is called to compute energies and gradients and the gradients for principal asymmetric unit are passed back to *phenix.refine* in place of conventional geometric restraints.

### S2. Refinement parameters

Parameters used in both sets of refinements.

~~~
c_beta_restraints=False
discard_psi_phi=False
strategy=*individual_sites individual_sites_real_space rigid_body
   *individual_adp group_adp tls *occupancies group_anomalous
flip_symmetric_amino_acids=True
refinement.target_weights.optimize_xyz_weight=True
refinement.main.number_of_macro_cycles=10
~~~

### S3. Full-dataset comparisons

Bond and angle rmsd comparisons (see figure S1) show that the bond rmsd values are numerically different but are smaller than the average sigma of 0.02Å (2pm) applied to protein bond restraints. Furthermore, the Amber angle rmsd values are approximately 2° across all resolutions – also lower than the average of ∼3° applied to protein angle restraints. The increased CDL/E&H rmsd values at high resolution may be result of the looser rmsd limit used for better than 2Å for the weight optimisation process. Comparing the means of the CDL/E&H and Amber rmsd values is not valid as force fields use more complex energetics rather than harmonic targets to ideal values.

### S4. Response to Bad Peptide Orientations

#### S4.1. Background

The low-resolution analysis of Cβ deviations in the main text made use of comparing the 1xgo structure at 3.5Å (Tahirov 1998) versus 1xgs at 1.75Å from the same paper. All six Cβ deviations in the Amber results versus none from CDL/E&H were compared, finding that in each case that Cβd was flagging an underlying problem: either a misfit side chain or an incompatibility between backbone and side chain.

For the issue of bad peptide orientations, however, only one example was illustrated (Figure 9). These problems are common at resolutions worse than 2.5Å, because the backbone CO direction is no longer seen (Richardson *et al*., 2018). Misoriented peptides can be diagnosed by CaBLAM (Williams 2018). CaBLAM uses virtual dihedral angles of successive Cαs and of successive COs to test whether the orientations of successive CO groups are compatible with the surrounding Cα trace. It flags outliers graphically in magenta on the CO-CO virtual dihedral. Since typically there is an energy barrier between widely different peptide orientations, the presumption is that refinement cannot easily correct these cases. However, that presumption needs to be tested.

#### S4.2. Most are not correctable by refinement

Ten cases were identified in 1xgo, for isolated single or double CaBLAM outliers (usually with other outliers also), surrounded by correct structure as judged in the same molecule at 1.75Å resolution (1xgs). For 6 of those 10 cases, neither CDL/E&H nor Amber refinement corrected the problem (His62, Thr70, Gly163, Gly193, Ala217, Glu286).

For example, figure S2 shows stereo images of the Glu286-Lys287 hairpin-loop case, where the CaBLAM outlier in 1xgo is accompanied by clashes, Ramachandran and rotamer outliers. Both CDL/E&H and Amber conformations are essentially identical to the original 1xgo, with no peptide improvement. They both remove all the clashes (clusters of hotpink spikes) and remove one of the six side chain outliers (gold) but not into the correct rotamer. In contrast, the high-resolution 1xgs, with very clear electron density (bottom panel), shows the Lys Cα and the two peptide carbonyl oxygens (red balls) differently placed by large distances and dihedral angles, forming a well H-bonded β**-**hairpin with no outliers of any kind.

#### S4.3. Other Outliers Often Better

In two cases the CDL/E&H results had fewer other outliers than Amber, although it did not actually reorient the peptide CO (Gly163, Gly193). The Gly163 case is shown in stereo in figure S3, for an S-shaped loop between non-adjacent β**-**strands, with two CaBLAM flags (magenta) and many other outliers. Both refinements remove the clashes, one of the rotamer outliers and one of the Ramachandran outliers (green). The CDL/E&H results in addition removed one of the CaBLAM outliers and the Cα-geometry outlier (red). However, neither refinement could manage the large rotation needed to correct the 163-164 peptide orientation, as judged by the more convincing conformation of the high-resolution 1xgs at bottom.

#### S4.4. Amber Sometimes Corrects Well

In three cases Amber managed a complete fix, while in contrast CDL/E&H did not improve (Asp88, Gly125, Pro266). The Asp88-Gly89 tight turn example is shown in Figure 9 of the main text.

Here in figure S4, the Gly125 loop example in a helix-helix connection is shown in stereo, to allow clear visualization of the CO orientation changes. 1xgo residues 121-126 (figure S3a) have two CaBLAM outliers (magenta dihedral lines) unchanged by CDL/E&H refinement (panel b). However, Amber refinement (panel c) manages to shift several CO orientations by up to 80° (red balls), enough to fix the CaBLAM outliers and to match extremely closely the better backbone conformation of 1xgs (panel d).

#### S4.5. A Partial Correction, Unconverged

Finally, in one especially interesting case (Lys22, in Figure S5a for 1xgo) Amber turned the CO (red circles) about halfway up to where it should be (panels b vs c), while CDL/E&H made no improvement to the peptide. The Amber model eliminated the Ramachandran and one of the CaBLAM outliers, but still had geometry outliers (a bond angle and a Cβ deviation). It seemed likely that Amber refinement had not fully converged and might move the CO all the way if run longer. A 30-cycle Amber run had earlier been done for 1xgo, without any major changes noticed beyond the 10-cycle. From that endpoint, two further runs were done, first of 30 cycles (“Amber60”), then a further 10 cycles (“Amber70”).

Figure S5d shows the fan of CO positions for all 7 of the deposits and refinements, progressively rotating counterclockwise from 1xgo to 1xgs. Indeed, both Amber60 and Amber70 successfully rotated the Lys22 peptide almost all the way to the good helical position seen in the high-resolution 1xsg (panel e), eliminating both the CaBLAM outlier and the intermediate-stage bond-angle outliers, presumably having crossed an energy barrier in the process.

One other CaBLAM-outlier peptide was corrected in Amber70 as well (Thr71). But for the Ala217 outlier, the wrong peptide was rotated, seduced by H-bonding to an Arg side chain in the wrong position.

In these long refinements, both R-factors and match to electron density suffer somewhat. In the cases examined, this often seems due to incorrect side chain rotamers (almost never correctable by refinement) pushing an otherwise-good backbone conformation a bit out of density (translated upward, for 1xgo Lys22). Future work will try to guide early correction of as many problems as feasible, for the faster and more successful refinement afterward that we now know is possible.

#### S4.6. Discussion

In summary, it is indeed true that refinement cannot usually correct a peptide orientation that is off by a large amount. The very tight geometry restraints in the CDL/E&H system presumably raise the barriers to peptide rotation. Amber is rather better at that, and about 1/3 of the time managed a good correction, although convergence can be very slow for such large changes. We feel it is crucial to try correcting problems such as flipped peptides in the initial model before refining it, however, crosstalk between backbone and side chains further complicates that process. However, we are enthusiastic about use of the Amber target to realistically improve conformation and especially sterics, once the model is mostly in the right local minima.

### S5. Boxplots of MolProbity results

Boxplots of the MolProbity results are given in figure S6 and S7. The latter has the difference between the CDL/E&H refinements and the Amber refinements. The largely (approximately 70% of all comparison) positive values show that Amber lowers the MolProbity score in the majority of cases.

---

**Figure S1.**
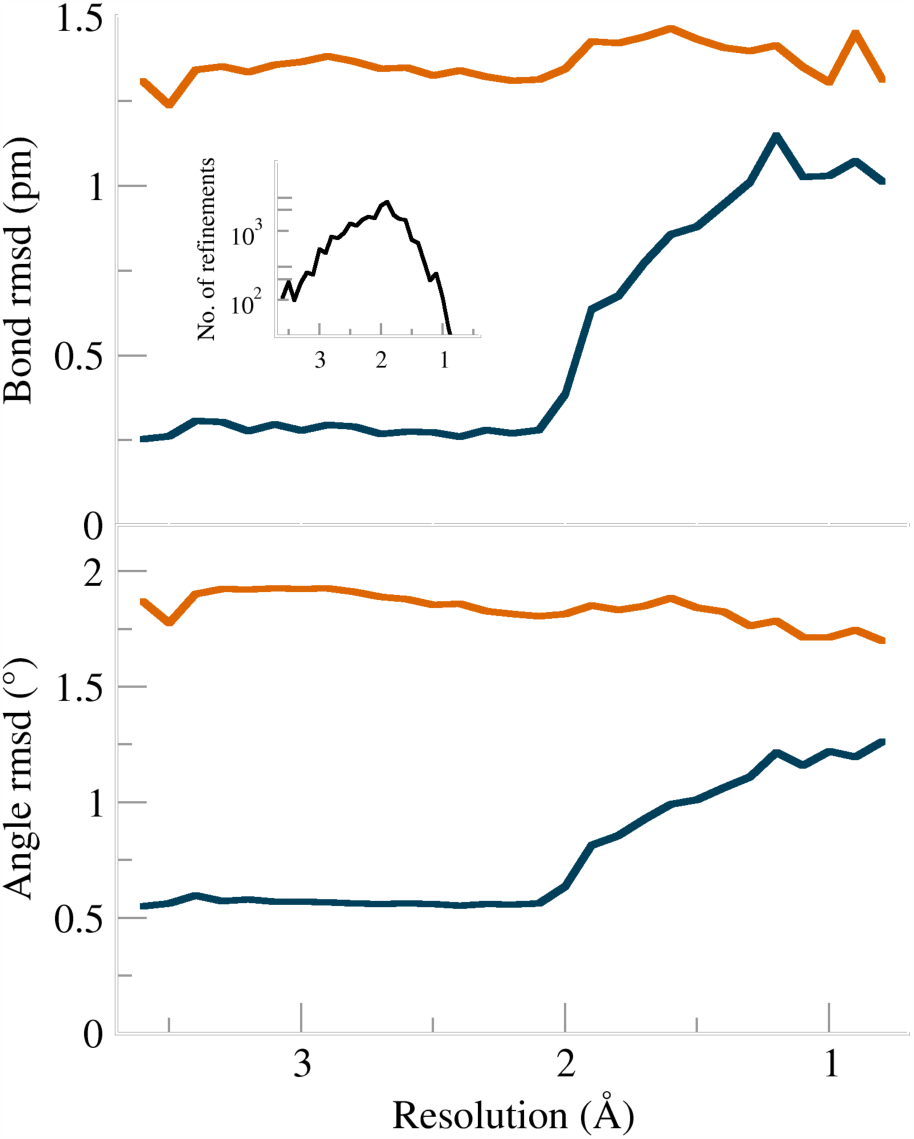
Bond and angle rmsd values for CDL/E&H (dark blue) and Amber (burnt orange) plotted against resolution.

**Figure S2.**
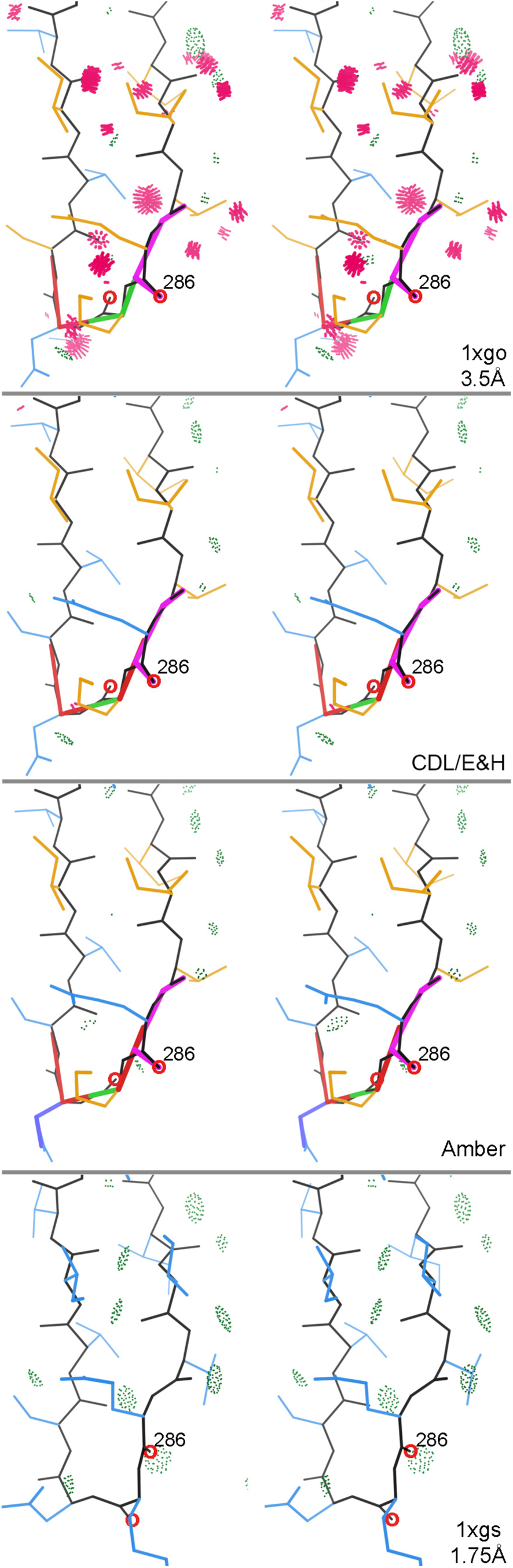
Stereo images of uncorrected CaBLAM problems for the beta-hairpin loop at Glu 286 - Lys 287 in 1xgo at 3.5Å resolution. a) As deposited, with outliers for CaBLAM (magenta lines on the CO dihedral), CaBLAM Cα-geometry (red lines on Cα trace), Ramachandran (green lines along backbone), rotamer (gold sidechains), and all-atom clash (clusters of hot-pink spikes) evaluations. b) As refined by Phenix CDL/E&H and c) as refined by Phenix Amber, both of which remove the clashes but do not correct the underlying conformation. d) In the 1xgs structure at 1.75Å resolution, showing a classic, outlier-free beta hairpin conformation with good backbone H-bonding and substantial corrections in peptide orientation and sidechain placement. The 286 and 287 peptide oxygens that move most are circled in red.

**Figure S3.**
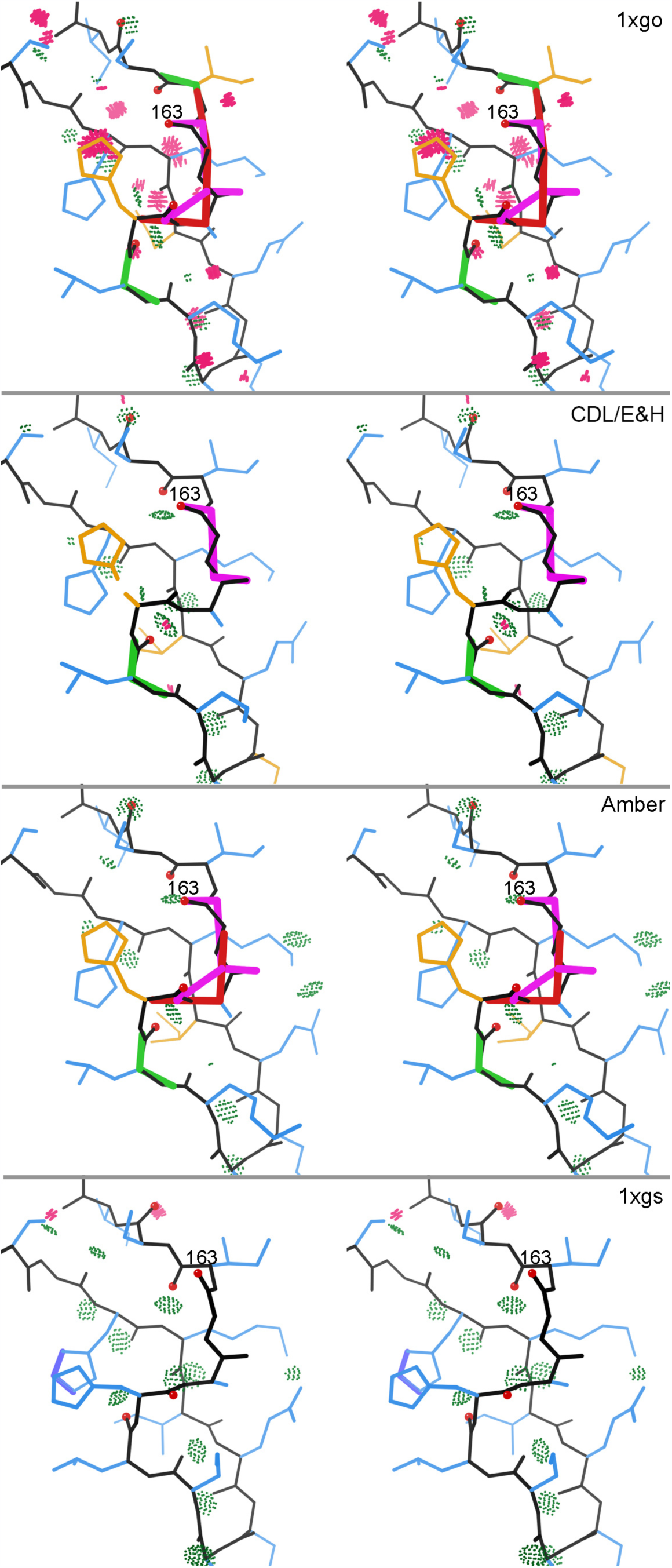
Partial correction of an S-shaped loop at 159-164 in 1xgo. a) As deposited, with many types of outliers. b) CDL/E&H corrects all but two backbone outliers. c) Amber corrects all clashes but few other outliers, and neither refinement changes the poor underlying conformation. d) The 1xgs structure achieves an outlier-free, well H-bonded conformation by shifting 4 peptide orientations (red ball on carbonyl O atoms), especially at Gly 163.

**Figure S4.**
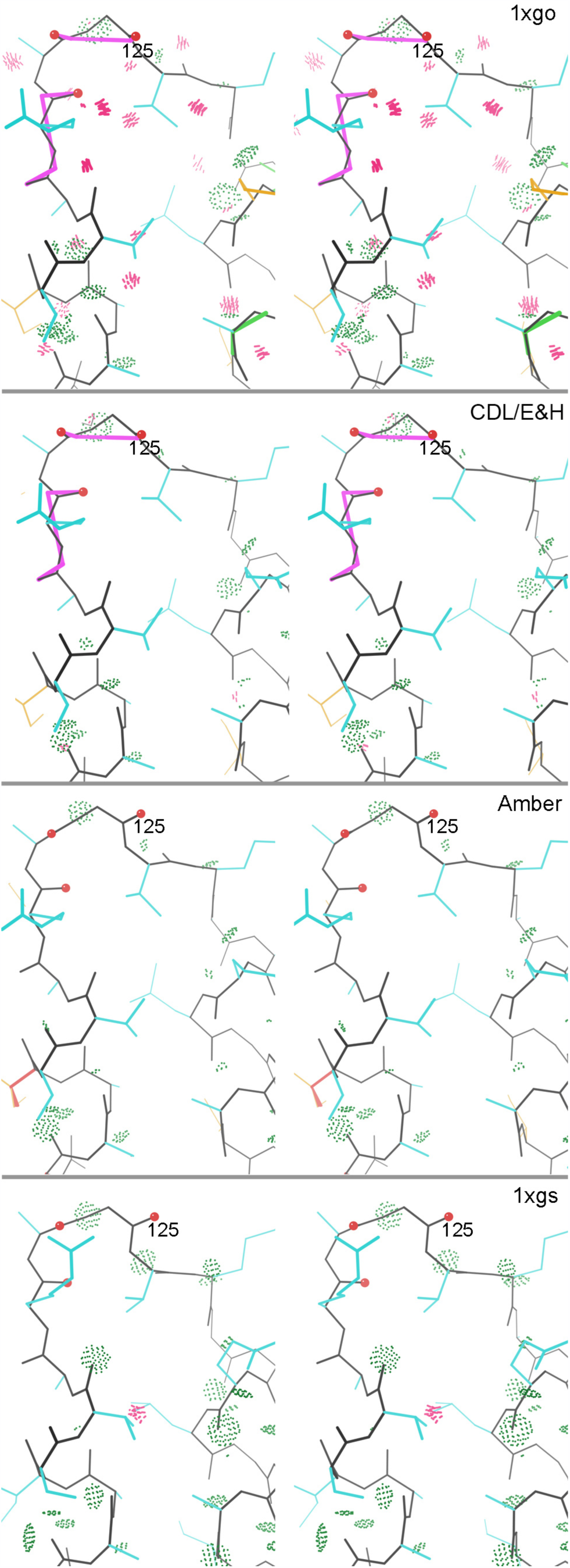
Successful Amber CaBLAM corrections in the helix-helix loop at 1xgo 121-126. a) As deposited, with clashes and two CaBLAM outliers. a) CDL/E&H corrects the clashes but not the backbone conformation. b) Amber reorients 3 successive peptides (red balls on peptide Os) by up to 80°, removing both CaBLAM outliers and matching extremely closely the conformation seen at high resolution in panel d.

**Figure S5.**
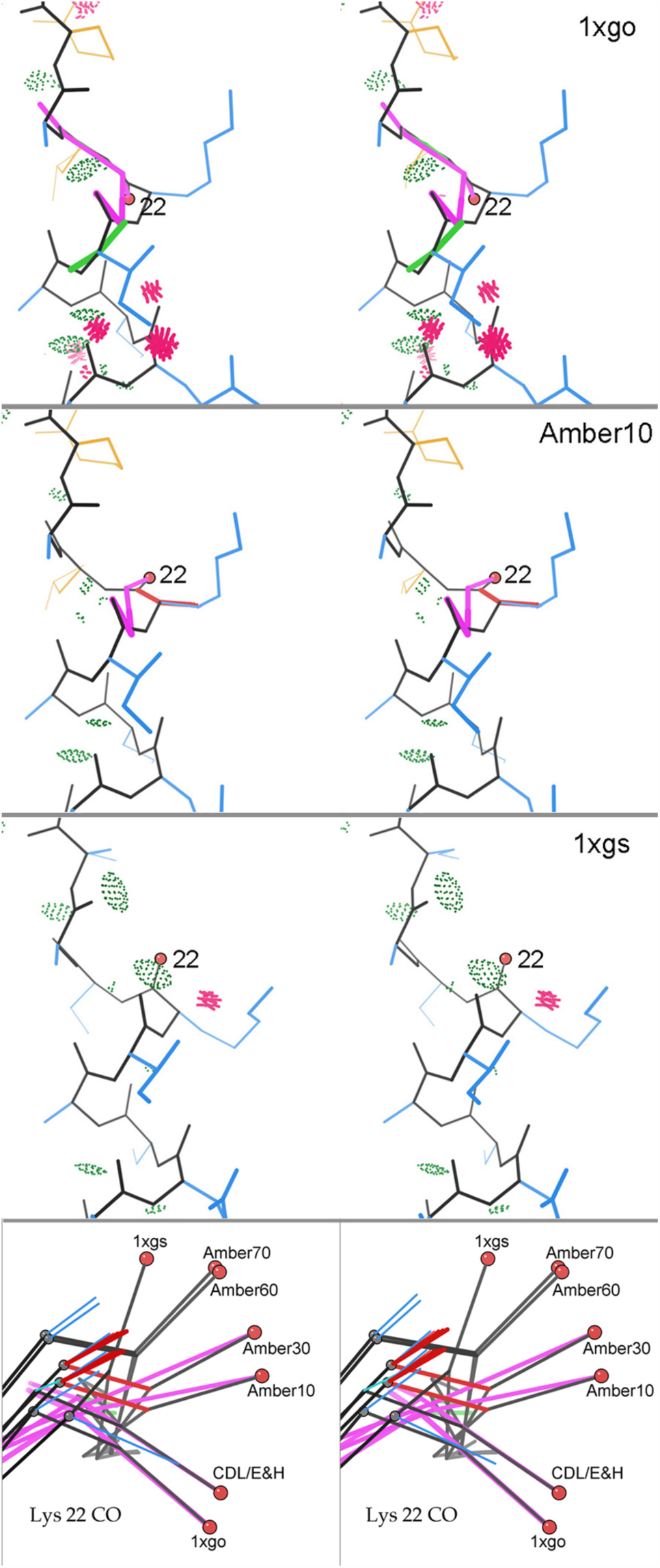
Gradual correction of the helix C-cap at 1xgo Lys 22. a) As deposited, with double CaBLAM outliers, clashes, and Ramachandran outlier. CDL/E&H refinement fixes clashes but leaves conformation unchanged. b) Amber refinement moves the crucial Lys 22 CO partway up toward α-helical orientation, relieving one of the CaBLAM outiers. c) Helical, outlier-free conformation of the C-cap region in 1xgs at high resolution. d) Superposition in side view, showing all Lys 22 CO orientations between 1xgo outlier and 1xgs α-helical: longer Amber refinement progressively corrects the orientation, converging close to the 1xgs orientation although with a translational shift we believe is an effect of incorrect sidechain rotamers.

**Figure S6.**
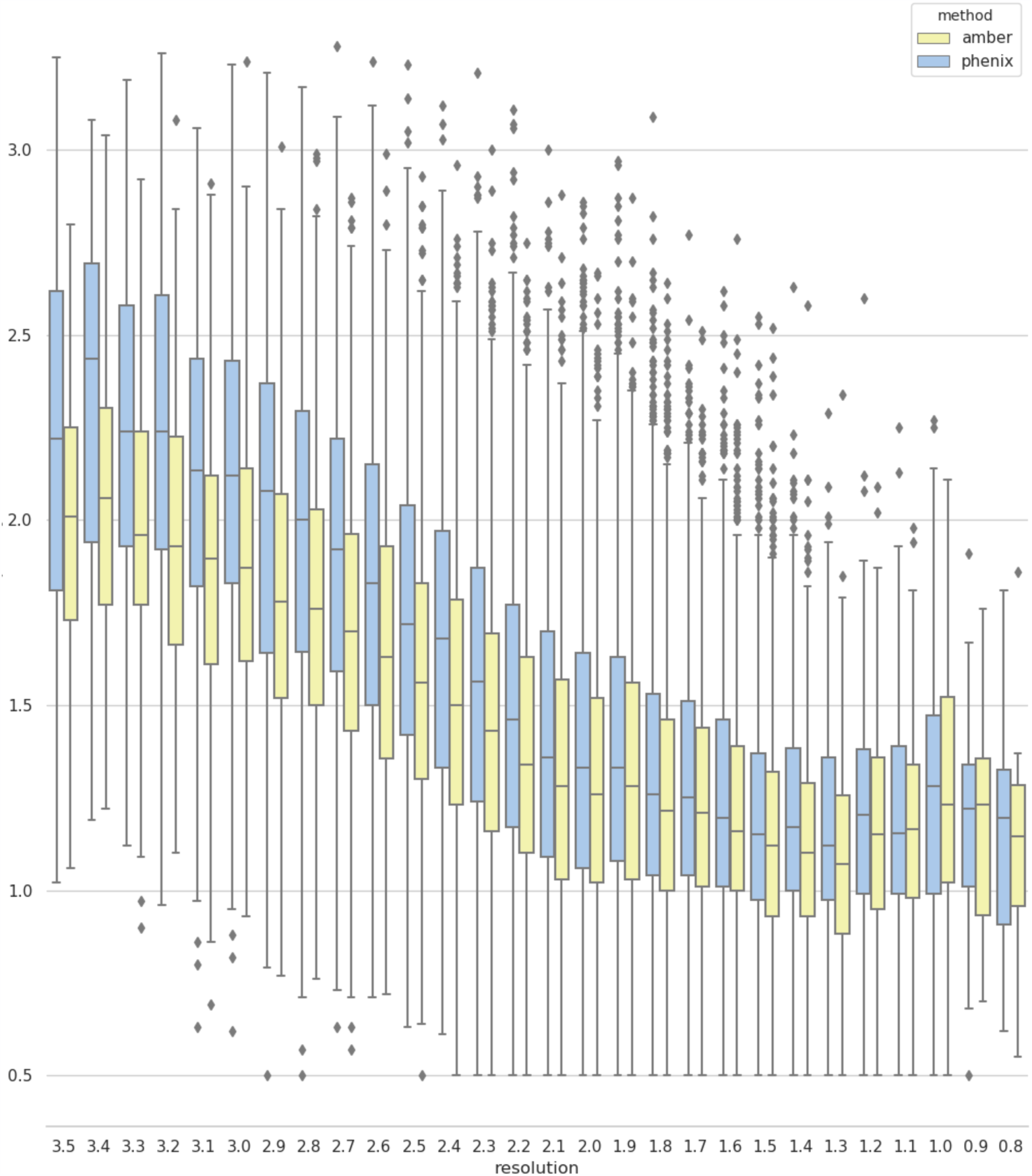
Molprobity scores binned into 0.1Å bins and plotted as boxplots. The box portion of the plot indicates the three quartile values while the whiskers cover the range of the values. Dots are outliers determined via a method of inter-quartile range.

**Figure S7.**
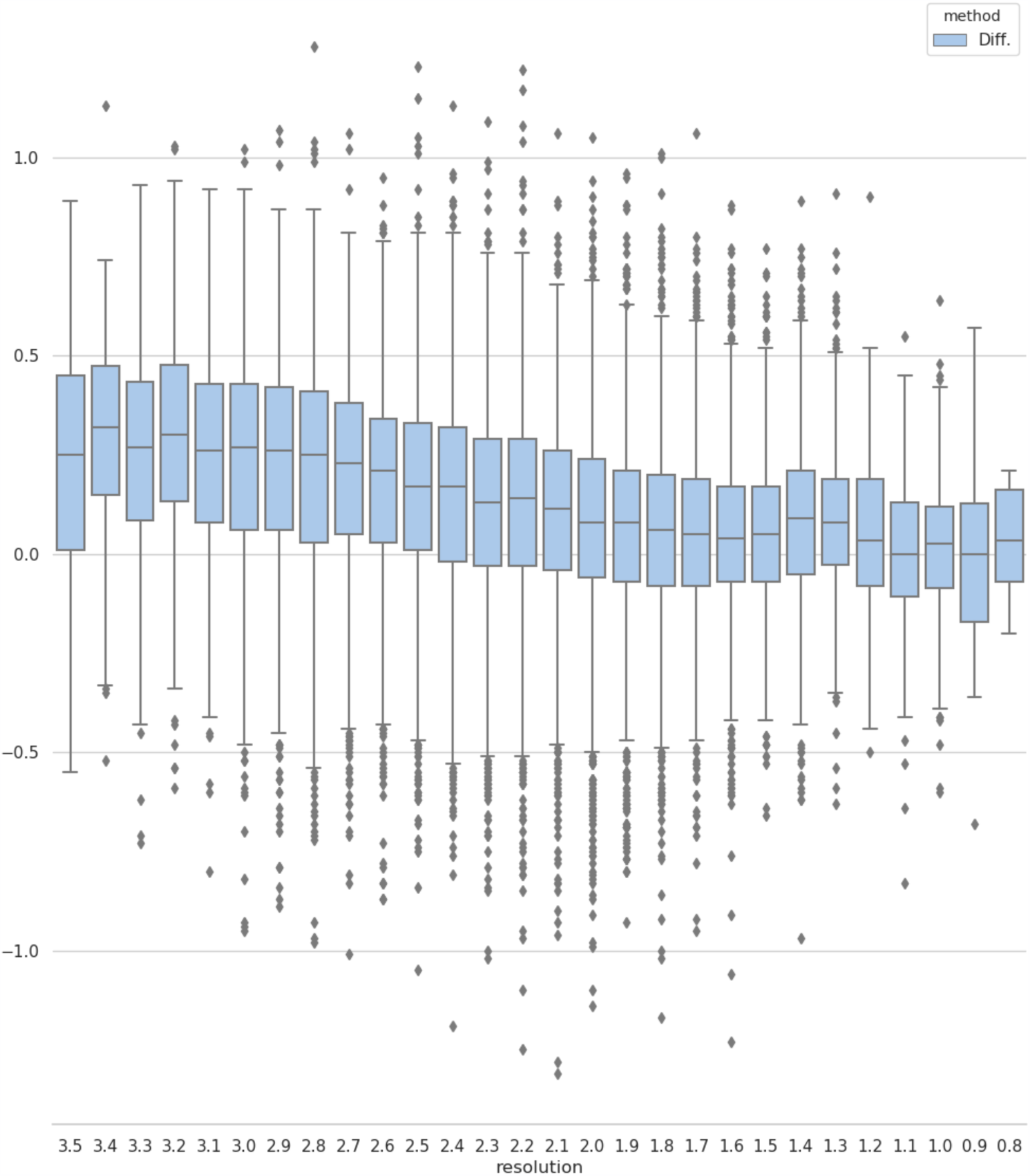
Differences in MolProbity scores between the Phenix and Phenix-Amber values (see figure S6) binned into 0.1Å bins and plotted as boxplots. Positive values indicate that Phenix-Amber refinements improved (decreased) the MolProbity score. The box portion of the plot indicates the three quartile values while the whiskers cover the range of the values. Dots are outliers determined via a method of inter-quartile range.

## Notes

#### Summary of Updates

Based on reviewer's comments. Most significant change to manuscript is in the supplemental material with in inclusion of boxplots for MolProbity scores.

